# Within-cycle instantaneous frequency profiles report oscillatory waveform dynamics

**DOI:** 10.1101/2021.04.12.439547

**Authors:** Andrew J. Quinn, Vítor Lopes-dos-Santos, Norden Huang, Wei-Kuang Liang, Chi-Hung Juan, Jia-Rong Yeh, Anna C. Nobre, David Dupret, Mark W. Woolrich

**Affiliations:** Oxford Centre for Human Brain Activity, Wellcome Centre for Integrative Neuroimaging, Department of Psychiatry, University of Oxford, OX3 7JX. UK; Medical Research Council Brain Network Dynamics Unit, Nuffield Department of Clinical Neurosciences, University of Oxford, Oxford, OX1 3TH, UK; Data Analysis and Application Laboratory, Innovation Centre, The First Institute of Oceanography, Qingdao, China; Pilot National Laboratory for Marine Science and Technology, Qingdao,China; Cognitive Intelligence and Precision Healthcare Centre, National Central University, Taiwan; Institute of Cognitive Neuroscience, National Central University, Taoyuan City, Taiwan; Department of Experimental Psychology, University of Oxford, Oxford. OX2 6GG. UK

## Abstract

Non-sinusoidal waveform is emerging as an important feature of neuronal oscillations. However, the role of single cycle shape dynamics in rapidly unfolding brain activity remains unclear. Here, we develop an analytical framework that isolates oscillatory signals from time-series using masked Empirical Mode Decomposition to quantify dynamical changes in the shape of individual cycles (along with amplitude, frequency and phase) using instantaneous frequency. We show how phase-alignment, a process of projecting cycles into a regularly sampled phase-grid space, makes it possible to compare cycles of different durations and shapes. ‘Normalised shapes’ can then be constructed with high temporal detail whilst accounting for differences in both duration and amplitude. We find that the instantaneous frequency tracks non-sinusoidal shapes in both simulated and real data. Notably, in local field potential recordings of mouse hippocampal CA1, we find that theta oscillations have a stereotyped slow-descending slope in the cycle-wise average, yet exhibiting high variability on a cycle-by-cycle basis. We show how Principal Components Analysis allows identification of motifs of theta cycle waveform that have distinct associations to cycle amplitude, cycle duration and animal movement speed. By allowing investigation into oscillation shape at high temporal resolution, this analytical framework will open new lines of enquiry into how neuronal oscillations support moment-by-moment information processing and integration in brain networks.

## 1: Introduction

Frequency, phase and amplitude have long been reported as important features of neuronal oscillations with behavioural and electrophysiological relevance. Furthermore, neuronal oscillations show non-sinusoidal waveform shapes that span a wide range of spatial and temporal scales (Cole and Voytek, 2017). Whilst waveform shape is emerging as a fourth relevant feature of neuronal oscillations, many theories of neuronal oscillations currently assume sinusoidal waveforms. This might be due to the fact that characterising and quantifying non-sinusoidal waveforms remains a substantial analytic challenge (Amzica and Steriade, 1998; Cole and Voytek, 2017). To uncover the role of waveform dynamics in rapidly unfolding brain activity, there is a growing need for novel analysis methods that are able to characterise a wide range of waveform shape features at the single-cycle level.

Waveform shape related parameters, such as skewness or asymmetry, can be estimated from higher order Fourier spectra such as the bispectrum or bicoherence (Bartz et al., 2019; Elgar, 1987; Sheremet et al., 2016). These methods require relatively long data segments to have high frequency resolution, and therefore do not provide single-cycle estimates. Alternatively, a set of waveform features for individual cycles can be described by the relative durations of different quartiles of a cycle (Belluscio et al., 2012; Cole and Voytek, 2019; Trimper et al., 2014). Whilst this approach is tractable on single cycles, the extracted features must be defined *a priori* and are limited to the resolution of the selected cycle control points such as the extrema and zero-crossings.

The temporal dynamics in oscillatory frequency can be quantified for a given waveform by its instantaneous frequency computed from the differential of the signal’s instantaneous phase (Boashash, 1992; Huang et al., 2009). Such instantaneous frequency estimates have been used previously in electrophysiology to explore dynamics in oscillatory peak frequency at high temporal resolution (Cohen, 2014; Liang et al., 2005; Nelli et al., 2017; Rudrauf et al., 2006). Crucially, any non-sinusoidal waveform features in an oscillation will lead to within-cycle instantaneous frequency modulations in which the frequency of an oscillation changes from moment-to-moment within a single cycle (Huang et al., 1998). The degree of non-linearity of an oscillation is related to the total amount of within-cycle frequency modulation (Huang et al., 2014; Tsai et al., 2016; Wang et al., 2012; Yeh et al., 2020).

We introduce a novel approach which creates smooth waveform shape profiles that describe non-sinusoidal features in single cycles with high temporal detail. To this end, we first operationalise waveform shape as the profile of instantaneous frequency across the cycle’s instantaneous phase. We then identify when and how an ongoing cycle deviates from a sinusoidal waveform by identifying points in the cycle where instantaneous frequency departs from a flat profile. For example, a cycle with a wide peak has a relatively low instantaneous frequency around the peak, and a cycle with a fast-ascending edge will have a relatively high instantaneous frequency between the trough and peak. In order to allow between cycle comparisons, we also need to account for how different cycles of an oscillation will play out at different speeds, leading to differences in extrema timing and overall duration. To overcome these problems, we introduce the process of ‘phase-alignment’ which reregisters the instantaneous frequency profiles onto a regularly sampled set of points in phase. To obtain the instantaneous phase time course of each cycle, we use the Empirical Mode Decomposition (EMD). EMD decomposes the time-series of interest into its oscillatory modes (Intrinsic Mode Functions; IMFs) that retain the non-stationary and non-linear signal features.

We outline and validate our novel approach in simulated data before applying it to theta-band oscillations recorded in the local field potentials (LFPs) of the mouse hippocampal CA1 during active exploratory behaviour. The hippocampal theta rhythm has a characteristic non-sinusoidal waveform shape (Belluscio et al., 2012; Buzsáki et al., 1985; Siapas et al., 2005) that is modulated by movement (Sheremet et al., 2016) and changes in sleep or drug states (Buzsáki et al., 1985). Using EMD to identify the theta rhythm, we show that phase-aligned instantaneous frequency is able to robustly characterise a continuous waveform shape profile for hippocampal theta. The results confirm the stereotyped fast-ascending and slow-descending shape in the cycle-wise average. Additionally, it reveals but with high amounts of shape variation across single cycles of theta, which we describe using a set of data-driven shape ‘motifs’. Finally, we find that cycle-level shape motifs have differential associations with theta amplitude, theta cycle duration and mouse movement speed. Overall, we demonstrate that behaviourally relevant dynamics in single-cycle oscillatory waveforms can be accurately and intuitively explored with phase-aligned instantaneous frequency profiles.

## 2: Methods

### 2.1: Data and Code Availability Statement

Code for the analyses in this paper are freely available online (https://github.com/OHBA-analysis/Quinn2021_Waveform) and data are available from the MRC BNDU Data Sharing Platform (https://data.mrc.ox.ac.uk/data-set/instantaneous-frequency-profiles-theta-cycles; requires free registration). The analyses in this study were carried out in python 3.7 using v0.4.0 of the EMD package (Quinn et al., 2021; https://emd.readthedocs.io/) and glmtools v0.1.0 for General Linear Model design and fitting (https://pypi.org/project/glmtools/). The wavelet transforms and principal components analysis were computed using SAILS v1.1.1 (Quinn and Hymers, 2020). The underlying python dependencies were numpy (Harris et al., 2020) and scipy (SciPy 1.0 Contributors et al., 2020) for computation and matplotlib (Hunter, 2007) for visualisation.

### 2.2: Masked Empirical Mode Methods

The EMD is implemented as a sifting algorithm that incrementally extracts the highest frequency features of a time-series into its oscillatory components known as IMFs (Huang et al., 1998). Once identified, the IMF is subtracted from the signal and the sifting process repeated to find the next fastest set of oscillatory dynamics. This process is iterated through until only a residual trend remains in the dataset, constituting the very slowest dynamics of the signal.

Transient or intermittent oscillatory signals can lead to a mix of different frequency components appearing in a single IMF; an issue known as mode mixing (Deering and Kaiser, 2005; Wu and Huang, 2009). To reduce mode mixing, we use an adapted version of the mask sift (Deering and Kaiser, 2005; Tsai et al., 2016). The mEMD involves the same core process as the original sift outlined above. However, at each iteration, a masking signal is added to the data before all the extrema (maxima and minima) in the masked signal are identified. An upper and lower amplitude envelope is then estimated by interpolating between the maxima and minima respectively. The average of the upper and lower envelope is subtracted from the data and the extrema identification, envelope interpolation and subtraction repeated until the average of the upper and lower envelopes are close to zero. The mask is then subtracted from the signal to return the IMF.

The performance of the sift is limited by a number of factors such as accuracy in peak detection and overshoot or edge effects in the envelope interpolation. As such, when applied to real data, the sift may not always perfectly isolate individual oscillations. To prevent envelope overshoot we used a monotonic PCHIP (Piecewise Cubic Hermite Interpolating Polynomial; implemented in scipy.interpolation.PChipInterpolator) interpolation during the sifting. The monotonic PCHIP interpolation avoids overshoot of the amplitude envelopes where the data are not completely smooth. When compared to a cubic-spline interpolation, this reduces instances where the envelopes become very large, or the upper and lower envelopes cross over.

#### 2.2.1: Frequency Transformation

The analytic form of each IMF was constructed using the Hilbert transform and the instantaneous phase set as the angle of the analytic form on the complex plane (See figure 2C for an example). To attenuate noise in the phase estimation, the unwrapped phase time-course was smoothed using a Savitsky-Golay filter (scipy.signal.savgol_filter; order = 1, window size = 3 samples). The instantaneous frequency (See figure 2D for an example) in Hertz is then computed from the derivative of this unwrapped phase:

**Figure 1:**
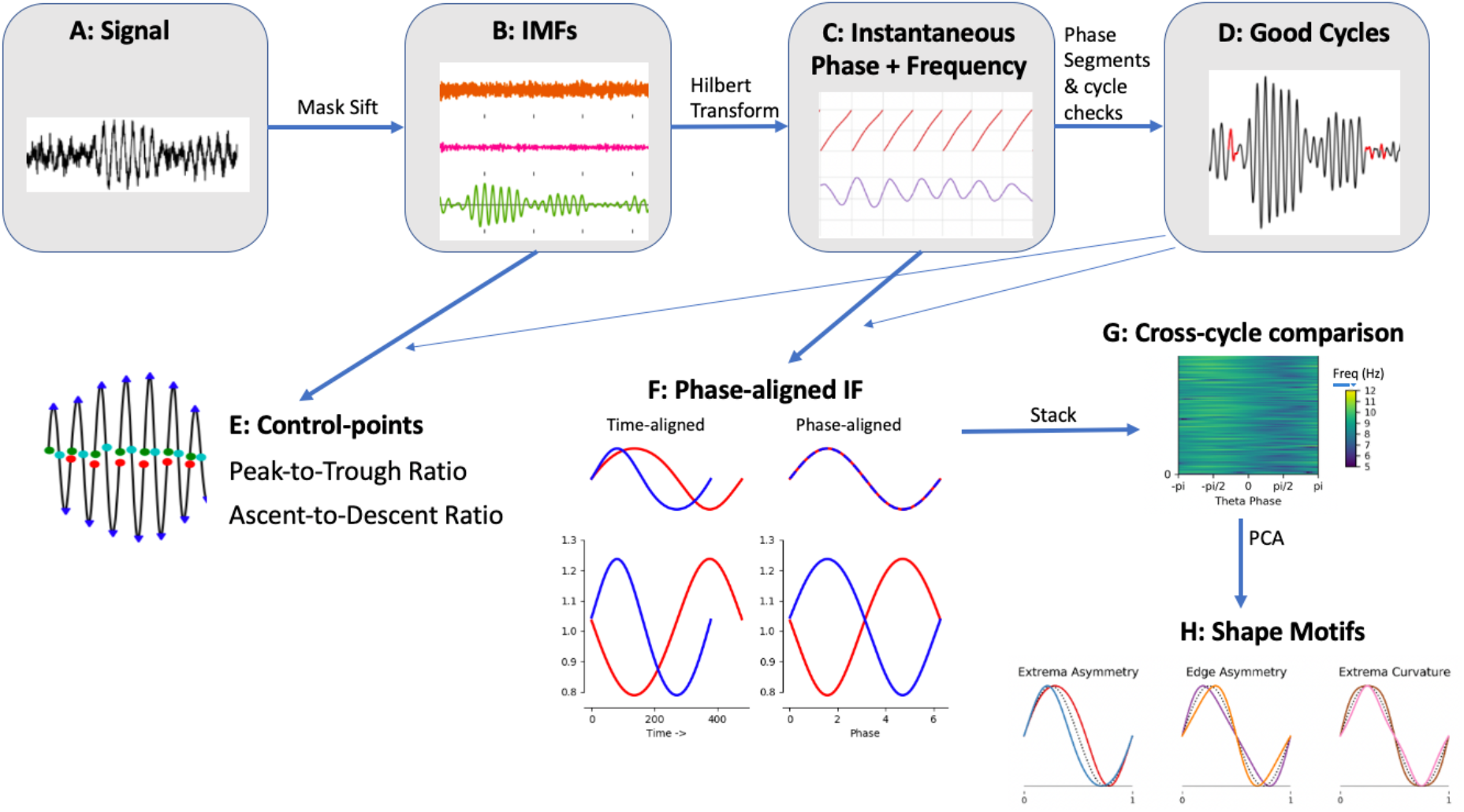
Overview of analysis applied to LFP recordings of hippocampal theta. A: The raw input LFP recording. B: The raw signal is split into Intrinsic Mode Functions (IMFs) using a mask sift. C: An instantaneous phase and instantaneous frequency time-course is estimated from the theta IMF using the Hilbert Transform. D: Cycle start and stop times are identified from jumps in the wrapped phase time-course and ‘bad’ cycles with distortions of reversals in phase are identified and removed. E: Control points (peaks, troughs, ascending zero crossings and descending zero crossings) are estimated from the good cycles within the theta IMF. Shape is then summarised using peak-to-trough and ascending-to-descending duration ratios. F: The instantaneous frequency of each good cycle is phase aligned to correct for variability in cycle duration and internal cycle timings. G: The phase-aligned cycles are stacked into a single array to allow for straightforward comparisons between cycles. H: A set of shape motifs are identified from the phase aligned IF using PCA.

**Figure 2:**
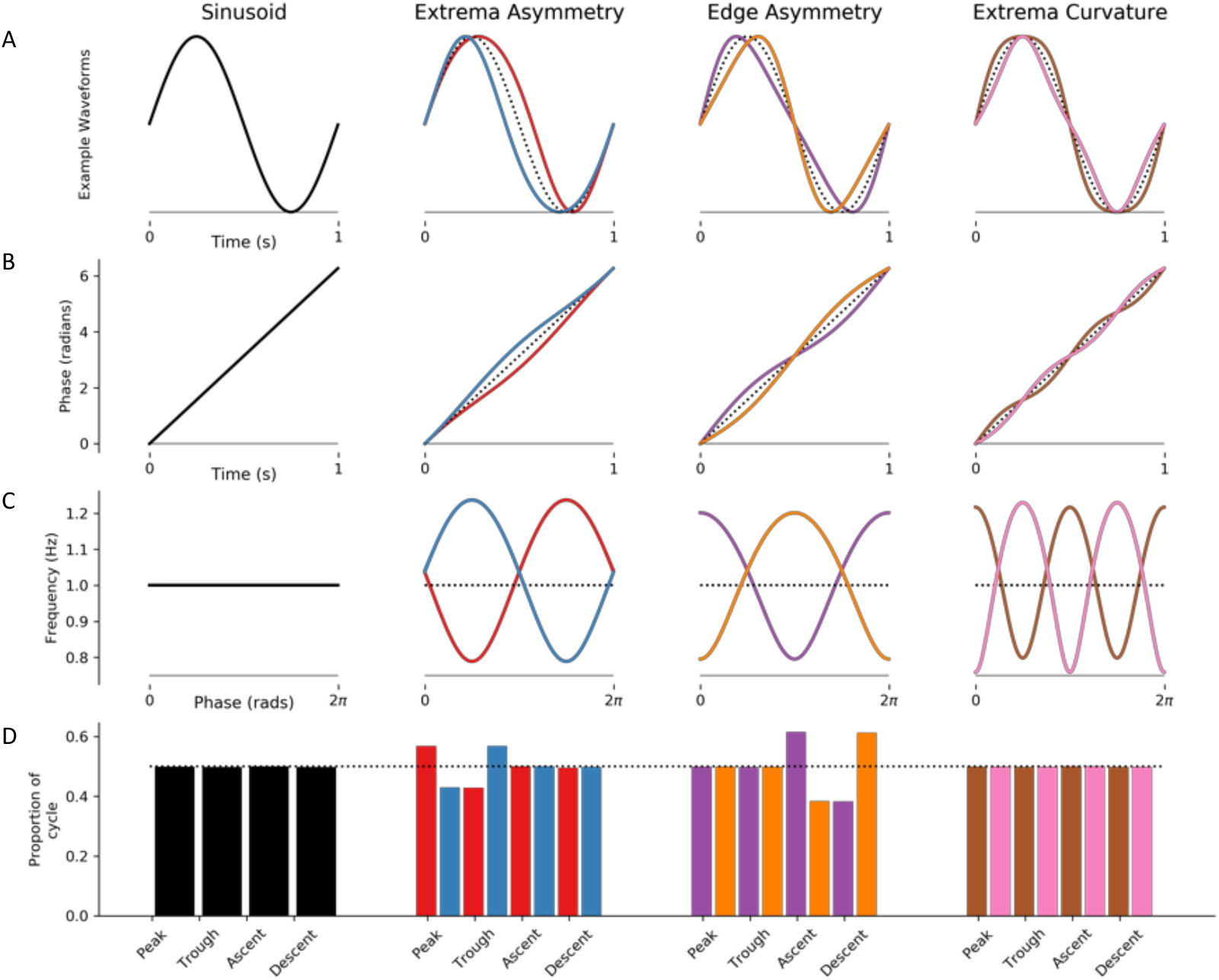
Instantaneous Frequency changes with oscillatory waveform shape. The four columns illustrate examples of different simulated oscillatory cycles with distinct waveform shapes. A: The time-domain waveforms for each cycle. The first column shows a sinusoid, and the remaining three columns show pairs of cycles with opposite waveform distortions (for reference a sinusoid is shown as a dotted back line). B: The instantaneous phase time course of the signals in the corresponding column. C: The instantaneous frequency time course of the signals in the corresponding column. D: The durations between different control points for each cycle the dotted line indicates the expected duration for a sinusoid.

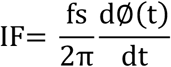

Where fs is the sampling frequency and ∅(*t*) is the unwrapped instantaneous phase time-course. The instantaneous amplitude time-course is computed as the absolute value of the analytic form of each IMF.

The distribution of instantaneous amplitude values by frequency or time and frequency can be computed from these instantaneous frequency metrics. A sparse matrix H**∈**R^(T × F) is filled with the instantaneous amplitudes value from the IMFs at their respective time and frequency co-ordinates. This matrix is the Hilbert-Huang Transform (HHT; See figure 2E for an example) and provides an alternative time-frequency transform to traditional Fourier based methods such as the short-time Fourier transform and the wavelet transform (Huang et al., 1998).

#### 2.2.2: Cycle Detection

The next stage is to segment the IMFs into their constituent cycles and identify which cycles will be included in further analysis. The start and end of theta cycles are located by the differentials greater than six in the phase. The start and end point of cycles in this paper is the ascending zero-crossing as this occurs at the point where the phase time-course wraps. Once identified, some cycles will be ‘bad’ in the sense that the oscillation captured by the IMF is not well represented, e.g., because the rhythm is not present over that time period or it is poorly estimated, and will be excluded from subsequent analyses. This is important for instantaneous frequency analyses as the differentiation step (see section 2.2.1) can be very noise sensitive. Included cycles are identified from the wrapped instantaneous phase time-course of the IMF containing the oscillation to be analysed. As the instantaneous phase computation via the Hilbert transform returns a value for every sample regardless of whether a prominent rhythm is present, only ‘good’ theta cycles are retained for further analysis. A good cycle is defined as having a phase with a strictly positive differential (i.e., no phase reversals) that starts with a value 0≤x≤π/24 and end within 2π-π/24≤x≤2π and 4 control points (peak, trough, ascending edge and descending edge).

#### 2.2.3: Control Point analysis

One approach to quantify oscillatory waveforms is to base analyses around parts of the cycle which can be straightforwardly identified in all cycles (Belluscio et al., 2012; Cole and Voytek, 2019). For example, the peak, trough, ascending zero-crossing and descending zero-crossings of a cycle can define a set of control points describing the relative timing of inflections and mid-points in a cycle. In the present analyses, the extrema (peaks and troughs) are detected by finding zero-crossings in the differential of an IMF. This initial extrema estimate is limited by the sampling frequency of the dataset and is refined using parabolic interpolation (Rato et al., 2008). The zero-crossings are initially identified from sign changes in the IMF time-course and refined by linear interpolation. The ratio of temporal durations between these control points can describe large scale shape features. Finally, we compute the peak-to-trough ratio and the ascent-to-descent ratios for each cycle (Cole and Voytek, 2019).

#### 2.2.4: Phase-alignment

We present an alternative approach to control points that ensures that entire waveform profiles can be combined and/or compared across cycles despite cycle-by-cycle differences in progression rate and overall duration. To compare waveforms across cycles that play out at different speeds, we use phase-alignment to register cycles onto a common grid. Phase alignment is performed on the instantaneous phase of a cycle and a measure of interest, such as the instantaneous frequency, which is observed at the same time-intervals. A linear one-dimensional interpolation function is fitted between the instantaneous phase (as x values) and the instantaneous frequency (as y values). The interpolation function is evaluated on a template set of instantaneous phase values with a linear spacing between 0 and 2pi, if any points in the template fall outside the fitted range, the interpolator returns an extrapolated value. This interpolated version of instantaneous frequency is then directly comparable across cycles as each point in the phase will occur at the same time. We compute phase-alignment using a linear interpolation across 48 fixed points across the zero to 2pi phase range.

Once an instantaneous frequency profile has been phase aligned, we can visualise a normalised waveform by projecting the frequency content back to a phase-time course. This is achieved by re-normalising the instantaneous frequency from Hertz back to radians in order to create a profile of successive phase differences. The phase time-course is then reconstructed from the cumulative summation of these phase differences. An oscillatory waveform can then be computed by taking the sine transform of this phase time-course. The resulting waveform has an amplitude of one and a consistent time-axis for all cycles. This ‘normalised waveform’ allows for visualisation of shape between cycles with different durations and amplitudes.

#### 2.2.5: Describing shape an instantaneous frequency mean vector

A simplified summary of a cycle’s shape can be computed from a mean-vector of the phase aligned instantaneous frequency according to the following equation:

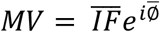

Where 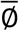 is the uniform phase-grid used in phase-alignment and 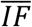 is the phase-aligned instantaneous frequency. This is similar to the mean-vector approach to computing phase-amplitude-coupling (Canolty et al., 2006). The mean-vector of a sinusoidal cycle will be zero whilst a non-sinusoidal cycle will return a complex value whose angle indicates which phase has the highest instantaneous frequency and the magnitude indicates the extent of the frequency modulation through the cycle. This method provides a straightforward summary but is only sensitive to unimodal deviations from a flat instantaneous frequency profile.

#### 2.2.6: Principal Components shape motifs

A more complete, data-driven approach to summarising shape from instantaneous frequency uses Principal Components Analysis (PCA) to identify the principal modes of variation in shape across the included cycles. Phase-aligned cycles are concatenated into a single matrix of size [nphases x ncycles]. The second dimension of this matrix is reduced to by PCA. This results in an [nphases x ncomponents] matrix of shape ‘motifs’ defined by the distribution of component weights across phase and an [ncomponents x ncycles] matrix of PC scores indicating the presence of each component motif in each individual cycle.

The component motif matrix defines the axes of variability in waveform shape across cycles. The shapes captured along each of these axes are visualised by defining a set of PC scores containing the maximum or minimum observed score for the PC in question and zeros for all others. These scores can be projected back into the original data space to provide exemplar instantaneous frequency profiles for both extremes of the PC axes. Finally, these exemplar IF profiles can be projected back into the time-domain to generate a normalised waveform that preserves the shape depicted in the exemplar IF profiles.

### 2.3: Simulation analyses

#### 2.3.1: Schematic cycle generation

To illustrate the relationship between waveform shape, instantaneous phase and frequency, a set of noise free oscillations were generated. First, a linearly progressing phase time-course is generated and sinusoid is created by taking a sine-transform of this wrapped phase. Different non-sinusoidal cycles are generated by modulating the unwrapped phase time-course by sine and cosine waves at different phases and frequencies. The cycles with extrema and edge asymmetry are generated by modulating the phase with a 1 Hz sine or cosine respectively.

The extrema curvature examples are generated by modulating the linear phase with a 2 Hz sinusoid. From the computed cycle time-courses the instantaneous phase and instantaneous frequency are re-estimated using the Hilbert transform. Finally, the waveform shape is represented by phase-aligning the instantaneous frequency time-course of each cycle type with its instantaneous phase.

#### 2.3.2: Noisy Signal Generation

A more realistic noisy simulation was used for the results in figures 3, 4 and 5. A simulated oscillation at 12 Hz was generated using an autoregressive oscillator with the following transfer function:

**Figure 3:**
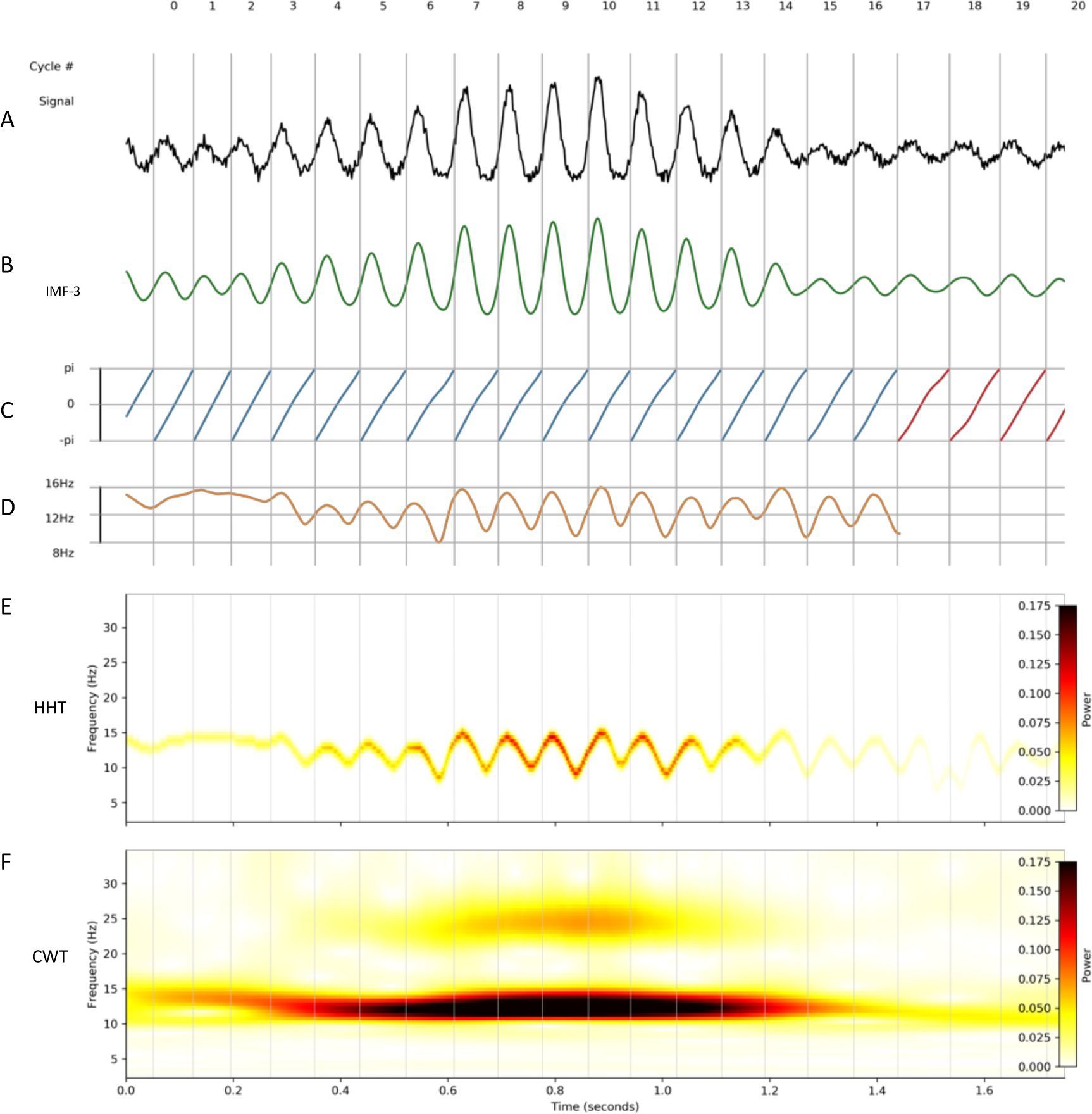
Instantaneous frequency analysis on a noisy non-sinusoidal oscillation. A simulated 12 Hz oscillation can be extracted from a noisy time-series and represented with EMD, instantaneous frequency and the Hilbert-Huang Transform. Vertical grey lines denote the starting times of individual cycles across the different panels. A: The simulated noisy non-sinusoidal oscillation. B: IMF-3 extracted from ‘A’ containing the simulated oscillation. C: Instantaneous phase time-course of IMF-3. Cycles excluded from further analysis are indicated in red. In this case, these cycles were below the amplitude threshold. D: Instantaneous frequency time-course of IMF-3. E: Hilbert-Huang Transform (HHT) of the simulated data segment. F: Continuous Wavelet Transform (CWT) of the simulated data segment.

**Figure 4:**
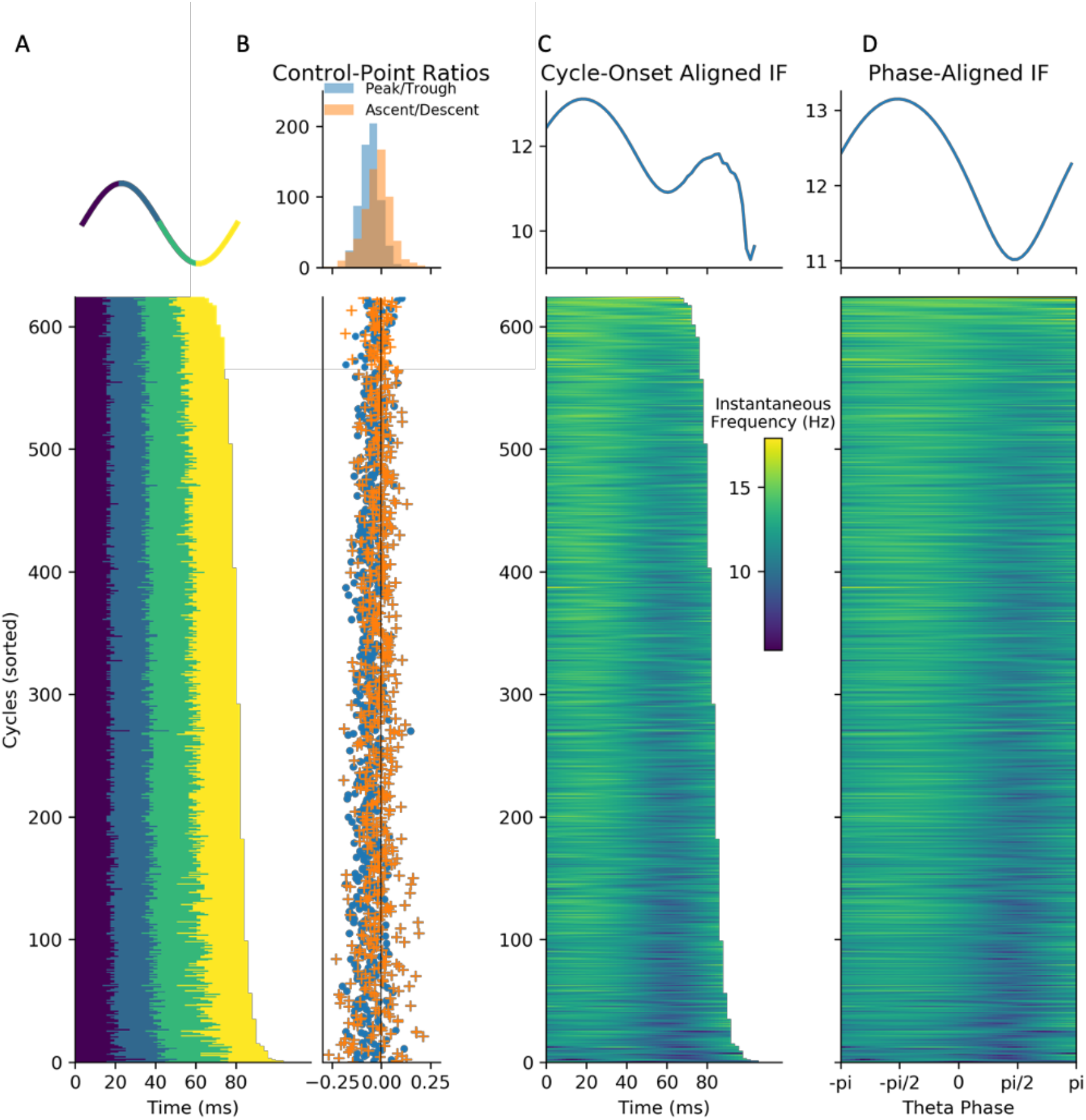
Methods for comparing waveform across cycles. Simulated data illustrating how waveform shape can be quantified using control point ratios, instantaneous frequency and phase-alignment. A : The durations between successive control points for each simulated cycle. B : Top: distributions of peak-to-trough and ascent-to-descent durations. Bottom: The peak to-trough and ascent-to-descent ratios for each cycle. C : Top: the average temporally aligned instantaneous frequency profiles. Bottom the temporally aligned instantaneous frequency profile for each cycle. D : Top: the average phase-aligned instantaneous frequency profiles. Bottom: the phase aligned instantaneous frequency profile for each cycle.

**Figure 5:**
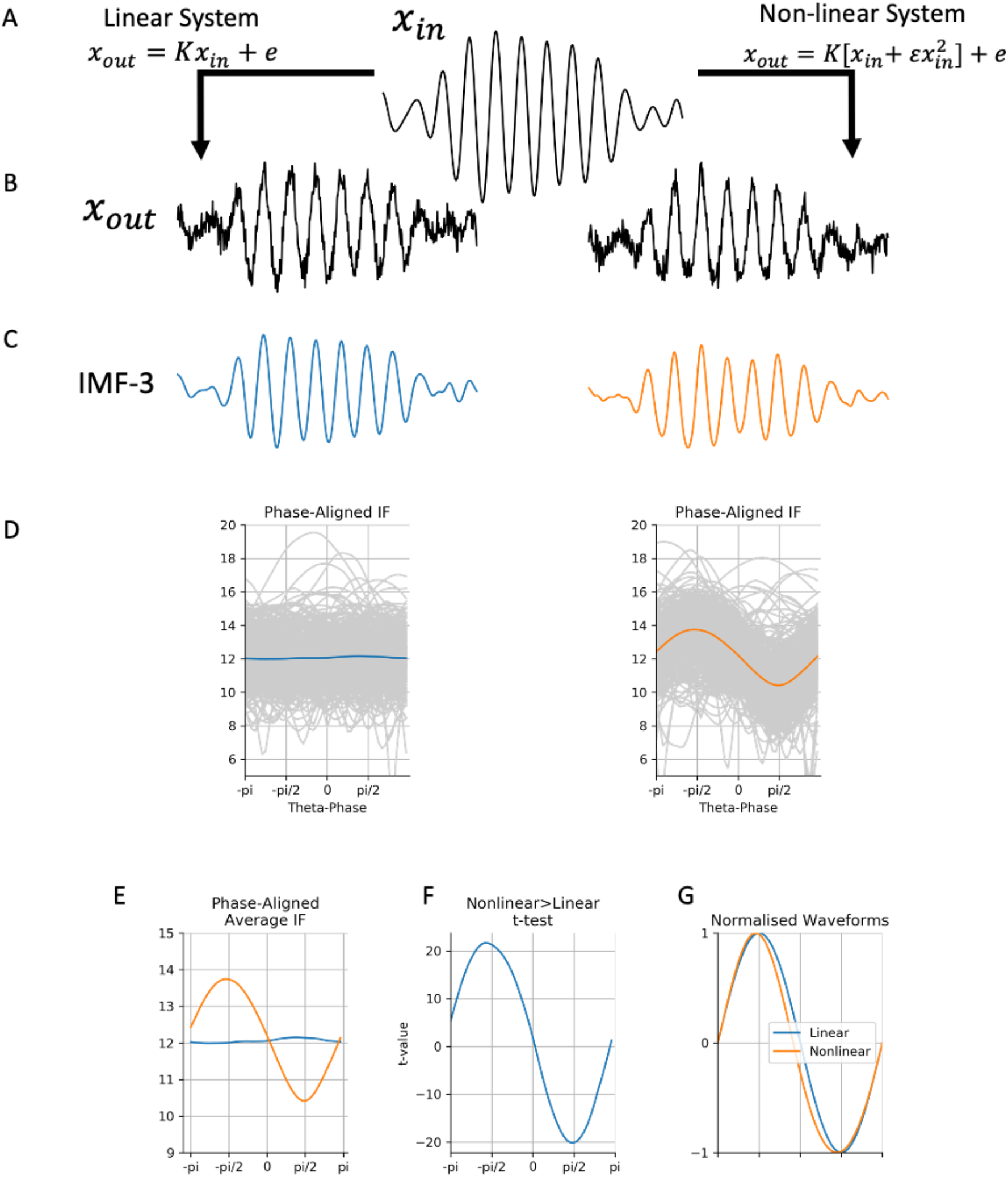
Comparing waveform shape in two simulated examples. A : A simulated oscillation is modulated by either a linear or a non-linear system. B : Output oscillations of the two systems. C : Oscillations recovered from the noisy simulation using EMD. D : Average phase-aligned instantaneous frequency profiles for the two systems with individual cycles in grey. E : Average instantaneous frequency profiles from ‘D’ overlaid together. F : t-values for a contrast between the instantaneous frequency in the two systems for each point in phase. G: Normalised waveforms for the two average frequency profiles in ‘E’.

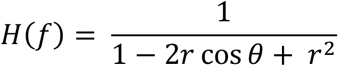

Where *θ* is the angular frequency of the oscillator (in rads/sec) and *r* is the magnitude of the roots of the polynomial (0 < r < 1). For this simulation, we computed *H* for r=0.95 and θ equivalent to 12 Hz and used its parameters to filter (a forwards and backwards filter using scipy.signal.filtfilt) white noise. This generates a noisy sinusoidal oscillation which contains random dynamics in the frequency and amplitude of each oscillatory cycle. Sixty seconds of data were generated at 512 Hz.

This simulated oscillation was then modulated by one of two equations defined in equations 50.24 and 50.25 in section 50-6 in Volume 1 of Feynman’s Lectures of Physics (Feynman et al., 2011). The first equation defines a linear system that scales the signal by a constant leaving the waveform shape unchanged.

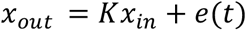

Whilst the second equation defines a nonlinear system that includes a term inducing a change in waveform shape as well as scaling the signal.

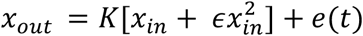

The nonlinearity in the second equation makes the peak of the oscillations shorter and widens the trough. Both systems include an additive white noise term.

#### 2.3.3: Noisy Signal Analysis

The simulations are separated into IMFs using the masked sift. The relatively straightforward dynamics in this simulation allow a simplified mask-sift to be applied. To define the mask frequencies, a first IMF is extracted using the standard sift routine. The number of zero-crossings in this IMF defines the frequency of the initial mask with subsequent masks being applied at half the frequency of the previous one. The mask amplitudes are set equivalent to 1-standard deviation of the previously extracted IMF.

The 12-Hz oscillation is isolated in the third IMF (IMF-3) of this mask sift. The frequency transformation of this IMF is computed using the Hilbert transform (section 2.2.1). The Hilbert-Huang transform (HHT) is computed using 64 frequency bins between 2 and 35 Hz. A wavelet transform was computed using a 5-cycle Morlet basis computed using the same 64 frequencies as the HHT.

The timing of individual oscillatory cycles is identified using the phase jumps in the instantaneous phase time course where the oscillatory amplitude was above a threshold of 0.04 and instantaneous frequency below 18Hz. We compute control-point ratios (section 2.2.3), time locked instantaneous frequency profile and phase aligned instantaneous frequency profile (section 2.2.4) across all included cycles (see section 2.2.2) of IMF-3.

The sift, frequency transform, cycle detection and shape metrics are computed for both the linear and non-linear systems defined in section 2.3.2 The difference between the linear and non-linear systems is quantified using a separate t-test on the peak-to-trough and ascending-to-descending control point ratios, and for each point in phase across the phase-aligned instantaneous frequency profiles. Finally, the normalised waveforms are computed from the average phase-aligned instantaneous frequency profiles from each system.

### 2.4: Hippocampal theta analyses

Local field potentials (LFPs) were recorded from the pyramidal layer of hippocampal CA1 using multi-channel tetrodes (Lopes-dos-Santos et al., 2018). Recordings were made during open-field exploration in both familiar and novel environments across six recordings taken over three recording days from each of three mice. Further data acquisition details can be found in appendix 1.

#### 2.4.1: Mask sift and Frequency transform

LFP recordings were each separated into oscillatory components using the mask sift (Figure 1 A & B) with masks placed at f_m = [350, 200, 70, 40, 30, 7, 1 Hz]. These masks were selected to capture components with frequencies above the mask frequency down to around f_m*0.7 (Fosso and Molinas, 2017; Rilling and Flandrin, 2008). Keeping the masks constant across recordings ensures that the frequency content of each IMF will be comparable across recordings. For instance, we isolate the theta oscillation in IMF-6 using a mask frequency of 7 Hz. These mask frequency parameters were validated by rerunning the phase-aligned instantaneous frequency analyses with jittered mask-frequencies (Supplemental section 8.2; figure S2), showing that the theta waveform description is robust to reasonable changes to the mask frequency values. These mask frequencies were effective in this set of CA1 LFP recordings, but it is expected that a different set of mask frequencies would be needed to analyse time-series containing different oscillatory dynamics. Next, a frequency transformation was computed for each IMF using the Hilbert transform and the methods from section 2.2.1 (Figure 1C).

#### 2.4.2: Cycle detection

To ensure that the detected theta cycles are physiologically interpretable theta activity, we identified cycles in each recording during times where the speed of movement of the mouse was greater than 1 cm/second. As faster movement is associated with stronger theta oscillations, this restriction increases the probability that our detected cycles represent physiologically interpretable theta events. We additionally restricted analyses to cycles in IMF-6 where cycle duration corresponded to 4-11 Hz frequency range (i.e., 312 and 113 samples respectively) and cycle amplitude was above the bottom 10% of the amplitude distribution. Finally, cycles that failed the cycle inclusion checks outlined in section 2.2.2 were removed from analysis at this point (Figure 1D).

#### 2.4.3: Cycle comparisons

We computed the temporally aligned instantaneous frequency profile, phase-aligned instantaneous frequency profile and normalised waveform for each included theta cycle. The average waveform shape within each dataset was estimated from the averaged phase aligned instantaneous frequency and a group average constructed from the mean of the six individual runs. Variability in waveform shape across single cycles in the group data is summarised using the instantaneous frequency mean vector (section 2.2.5) and visualised as a distribution in the complex plane in which the x-axis represents asymmetry between ascending and descending edge frequency and the y-axis represents asymmetry between peak and tough frequency. For comparison, we also identified the control points from each cycle of the theta IMF and constructed the peak-to-trough and ascending-to-descending duration ratios (Cole and Voytek, 2019).

#### 2.4.4: Waveform Motifs and Relation to behaviour

We next look to explore the waveform shapes which are present in the phase-aligned instantaneous frequency values. We use PCA (section 2.2.6) to identify the data-driven set of shape components that explain the most variance in the shape of theta cycles in this dataset. The first four principal components explaining 95% of variance defined our four shape motifs and were retained for further analysis. The reproducibility of the PCA is validated across 500 split-half iterations assessing the proportion of variance explained by each PC and the correspondence between the component shape in the two halves (Supplemental section 8.3; figure S3).

The relationship between the shape motifs and a set of three cycle covariates (i.e., cycle amplitude, cycle duration and mouse movement speed) was quantified using a General Linear Model (GLM). The GLM was created with a design matrix containing the mean and the three z-transformed covariates. These predictors were used to model the between cycle variability in the principal component (PC) scores for each shape component in turn. This resulted in four GLMs each fitting four parameter estimates. The t-statistic of each parameter estimate was computed, and statistical significance established using a row-shuffle non-parametric permutation scheme. 5000 permutations were computed for each PC-motif and dependent variable before statistical significance determined at p<0.01.

## 3: Results

### 3.1: Instantaneous Frequency tracks waveform shape

Figure 2 illustrates how instantaneous frequency reflects waveform shape in a set of noiseless simulated cycles (see section 2.3.1). A sinusoidal cycle (Figure 2A) has a monotonically progressing phase time-course which, in turn, has a flat instantaneous frequency profile. Analysis of the duration of different segments reveals that the peak, trough, ascending edge and descending edge all have the same duration. Cycles with a narrow peak, trough or descending edge show corresponding changes in their instantaneous frequency (Figure2 B & C). Specifically, the longer duration, slower features correspond to a lower instantaneous frequency. These instantaneous frequency profiles can describe a wide range of possible shapes. For example, cycles in which both the peak and trough are widened or pinched lead to instantaneous frequency profiles with multiple extrema (Figure 2: last column). While the simple control-point metrics used here can track individual waveform features such as peak or trough duration (Figure 2: bottom row), the quantification of more complex shapes would require the definition of additional control points and shape metrics.

### 3.2: Quantifying and comparing waveform shape in a simulated signal

We next use simulations to illustrate how instantaneous frequency analyses can be conducted on a noisy signal with a dynamic 12 Hz oscillation modified by a non-linearity which widens the trough of each cycle (see section 2.3.2). This oscillation was isolated from the noisy background using mask sift (Figure 3B) before the Hilbert transform was used to compute the instantaneous phase time-course (Figure 3C). It is evident that the phase time-courses do not progress linearly through all cycles; these deviations from monotonic phase progression are quantified in the instantaneous frequency time-course (Figure 3D). Instantaneous frequency sweeps within a single cycle reflect the non-sinusoidal shapes of the time-domain waveforms. For this simulation, the instantaneous frequency tends to be higher during the first half of the cycle and lower in the second half, reflecting the non-linearity that shortens the peak and widens the trough of these oscillations.

The HHT of the simulated signal (Figure 3E) retains the high time-frequency resolution of the instantaneous frequency time-course allowing within cycle frequency dynamics to be visible. In contrast, while a standard 5-cycle Morlet wavelet transform identifies similar power dynamics, variability in frequency within single cycles are not resolved (Figure 3F). A further disadvantage is that the non-sinusoidal waveform shape of this simulation introduces a 24Hz harmonic component into the wavelet transform.

Individual cycles of an oscillation play out at different rates leading to differences in the timing of extrema within cycles and in overall cycle duration. These two sources of variability hamper comparisons between individual oscillatory cycles. As outlined above (section 2.2.3) one method to solve this issue is to discretise the cycle using a set of control points before computing the proportion of time spent in different segments of the cycle (Belluscio et al., 2012; Cole and Voytek, 2019). The cycle phase quartiles (ascending zero-crossing, peak, descending zero-crossing and trough) of 500 cycles of the simulated signal is shown in Figure 4A. The ratios of peak-to-trough duration and ascending-to-descending time of these cycles suggests longer troughs and shorter peaks, whilst the ascending and descending portions of the cycle are approximately equal in duration (Figure 4B).

An alternative approach for comparing cycles is to align the instantaneous frequency profiles to one of the control points. For example, we aligned the 500 cycles to the ascending zero-crossing and computed their time-locked average (Figure 4C). The time-locked instantaneous frequency profile of these cycles is not flat, reflecting the presence of non-sinusoidal shape in this simulated signal. However, the precise type of non-sinusoidal shape is ambiguous from this average, due to variability in the location of different waveform features within single cycles. In this case, the instantaneous frequency is highest around 10 samples after the ascending zero-crossing; however, this time-lag might correspond to different points in the waveform in different cycles. In addition, variability in the duration of cycles means that, after a certain point, different numbers of cycles contribute to the average, making the estimate unstable.

Here we present an approach that overcomes these shortcomings. In brief, phase-alignment removes this ambiguity by visualising the instantaneous frequency of a cycle across a fixed grid of points along its phase (see section 2.2.3). For instance, an oscillatory peak is normalised to occur at the same phase value irrespective of the cycle’s duration or shape. This corresponds always to exactly one quarter of the phase of each cycle, but not necessarily to one quarter of the duration of each cycle. By aligning the instantaneous frequency to the phase, we remove the temporal distortions caused by varying shapes and cycle durations, and express the shape with the phase-aligned instantaneous frequency values. The phase-aligned instantaneous frequency of the simulated cycles (Figure 4D) now unambiguously shows the increased frequency around the peak of the 12Hz oscillation and decreased frequency around the trough. The average across the phase-aligned cycles is then a smooth representation of the shape of the entire cycle.

Comparisons between sets of cycles is straightforward once waveform shape has been estimated from instantaneous frequency and normalised through phase-alignment. This is illustrated by contrasting the shape of a noisy a 12-Hz oscillation modulated by either a linear or non-linear system (section 2.3.2; Figure 5A). The non-linear system has a wide-trough shape whilst the linear system has a sinusoidal waveform (Figure 5B & C). The average phase-aligned instantaneous frequency values for the linear system correspond to a flat line at 12 Hz throughout the cycle. In contrast the non-linear system has an increased instantaneous frequency in the first half of the cycle and a decreased frequency in the second half (Figure 5D). We can compare the average instantaneous frequency profiles from both systems (Figure 5E) and compute conventional t-statistics (Figure 5F) to quantify any differences in waveform shape. In this simulation, we find that the non-linear system creates oscillations with increased frequency peaks and decreased frequency troughs.

Finally, the phase-aligned average frequency profiles can be projected back into a ‘normalised waveform’ to more intuitively visualise the type of non-sinusoidal distortions. These normalised waveforms have a constant duration and an amplitude of one, but retain any distortions in waveform shape quantified in the instantaneous frequency profile. For this simulation, the normalised waveform reveals the pinched peak and the widened generated by the non-linear system (Figure 5G).

### 3.3: Characterising waveform shape in Hippocampal Theta

LFP data recorded from the mouse hippocampus were analysed to explore the utility of phase-aligned instantaneous frequency as a measure of waveform shape. Figure 6A shows a three-second LFP recording from the pyramidal layer of the mouse dorsal CA1 (black) overlaid with the EMD-extracted theta IMF (red, Figure 6B shows all IMFs). In this case, the theta oscillation was isolated into IMF 6 with minimal disruption to its amplitude or waveform shape dynamics. Many of the theta cycles within this window have prominent non-sinusoidal waveform shapes, which are qualitatively visible in both the raw data trace (Buzsáki et al., 1986, 1985) and the EMD-extracted theta IMF. Importantly, the oscillatory waveform shape varies between successive cycles, though the amplitude and duration of the theta cycles are relatively consistent.

**Figure 6:**
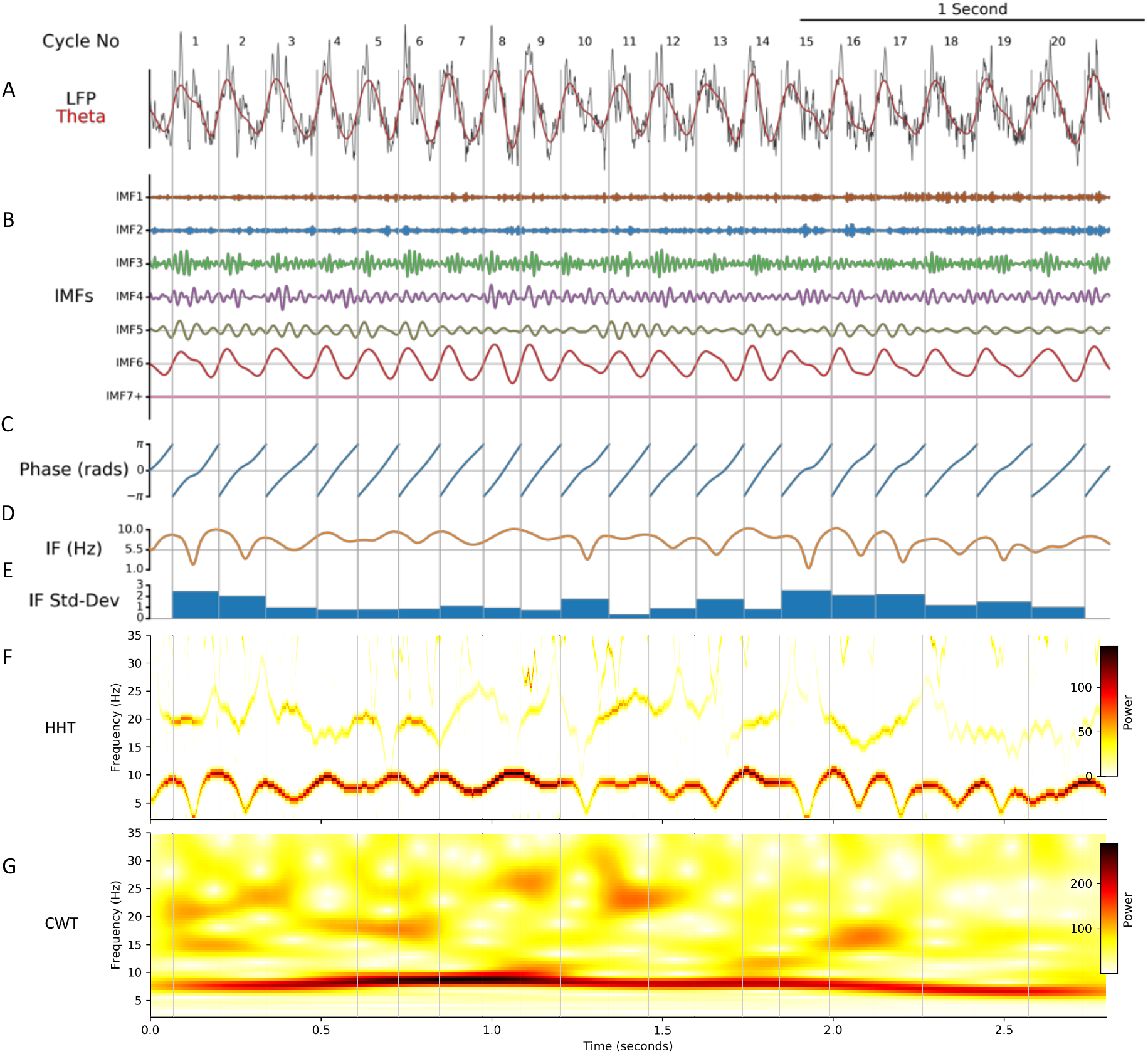
EMD analysis of a LFP segment containing hippocampal theta oscillations. A : A segment of a hippocampal LFP recording (black) overlaid with the extracted theta oscillation (red). B : IMFs extracted from this data segment using the mask-EMD. The theta oscillation is isolated into the IMF-6. C : Instantaneous phase time-course of the theta IMF. D : Instantaneous frequency time-course of the theta IMF. E : Variability in instantaneous frequency for each theta cycle. F : Hilbert-Huang Transform (HHT) of the LFP segment. G : Continuous Wavelet Transform (CWT) of the LFP segment.

The instantaneous phase (Figure 6C) and instantaneous frequency (Figure 6D) were computed from the theta oscillation in IMF-6. As with the simulation analysis, any within-cycle dynamics in the instantaneous frequency naturally represent the waveform of each cycle. This was summarised with the standard deviation of instantaneous frequency values within each cycle (Figure 6E). As an illustration, cycles 5, 9 and 11 have relatively sinusoidal shapes with flat instantaneous frequency profiles and low frequency variability. In contrast, cycle 13 is relatively non-sinusoidal with a dynamic instantaneous frequency profile and high frequency variability. The HHT provides a time-frequency description with sufficient resolution to depict these within cycle instantaneous frequency sweeps (Figure 6F). In contrast, a 5-cycle wavelet transform of the same data was not able to resolve these dynamics (Figure 6G).

Looking at individual cycles illustrates how instantaneous frequency can characterise waveform shape (Figure 7). Frequency increases and decreases correspond to slowing down and speeding up of the cycle as its waveform shows non-sinusoidal behaviour. It is evident that there are many observed shape profiles. For instance, the cycles labelled as *i* and *iii* in figure 7 had frequencies that dip during the centre of the cycle, indicating an elongated, low frequency descending edge. Cycle *v* had slowest frequency around -pi/2 corresponding to a wide peak. In contrast, cycle *vi* had relatively high frequency around -pi/2 and lower frequency at +pi/2 leading to a short, pinched peak and an elongated trough. Overall, the phase-aligned instantaneous frequency profiles and normalised waveforms provide a rich description of oscillatory waveform, despite wide variability in cycle amplitude, duration and shape.

**Figure 7:**
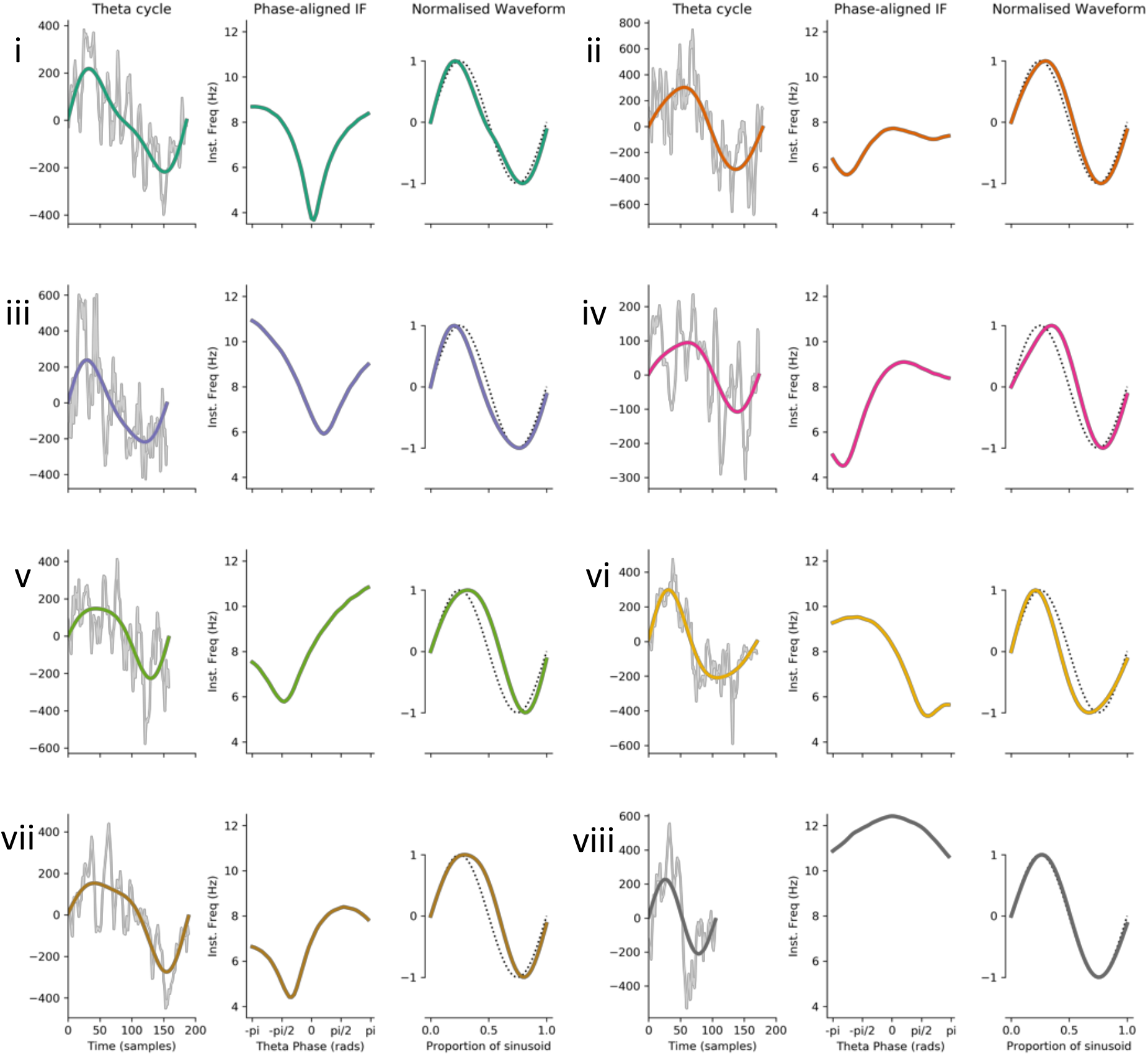
Characterising shape in eight example cycles of hippocampal theta. Eight representative theta cycles. For each example, the first subpanel shows the raw data (grey) with the theta IMF super imposed (coloured line). The second subpanel shows the phase aligned instantaneous frequency and third panel shows the normalised waveform (coloured line) with a sinusoid for reference (black dotted line).

### 3.4: Using phase-aligned instantaneous frequency to compare cycles

We next computed waveform shape from around 1500 hippocampal theta cycles from a single recording using three different methods: control-point ratios (see section 2.2.6), control-point locking, and phase-alignment (see section 2.2.3). The durations between specified control points were computed for each cycle (Figure 8A) and the peak-to-trough ratio and the ascent-to-descent ratio computed (Figure 8B). The peak-to-trough ratios are evenly distributed around zero, whereas there is a bias in the ascent-to-descent ratios suggesting that the descending edge of theta is longer than the ascending edge. The instantaneous frequency profiles locked to the ascending zero-crossing show a wide variety of shapes with a group average tendency for frequency to start around 9 Hz and to decrease through the duration of the cycle (Figure 8C). As described above, this average effect is challenging to interpret due to within-cycle variability in the timing of cycle features and between cycle variability in total cycle duration. Our proposed phase-aligned instantaneous frequency profiles (Figure 8D) resolve these ambiguities. This shows that theta cycle instantaneous frequency in this single recording starts around 9 Hz at the ascending zero-crossing, decreasing to around 8.1 Hz at the descending zero-crossing, before increasing again to 9 Hz at the end of the cycle. This is consistent with a fast-ascending and slow descending cycle shape revealed by the control point analysis and in previous literature. The phase-aligned instantaneous frequency approach is able to show this effect as a continuous shape profile for single cycles, which can be straightforwardly compared at the group level.

**Figure 8:**
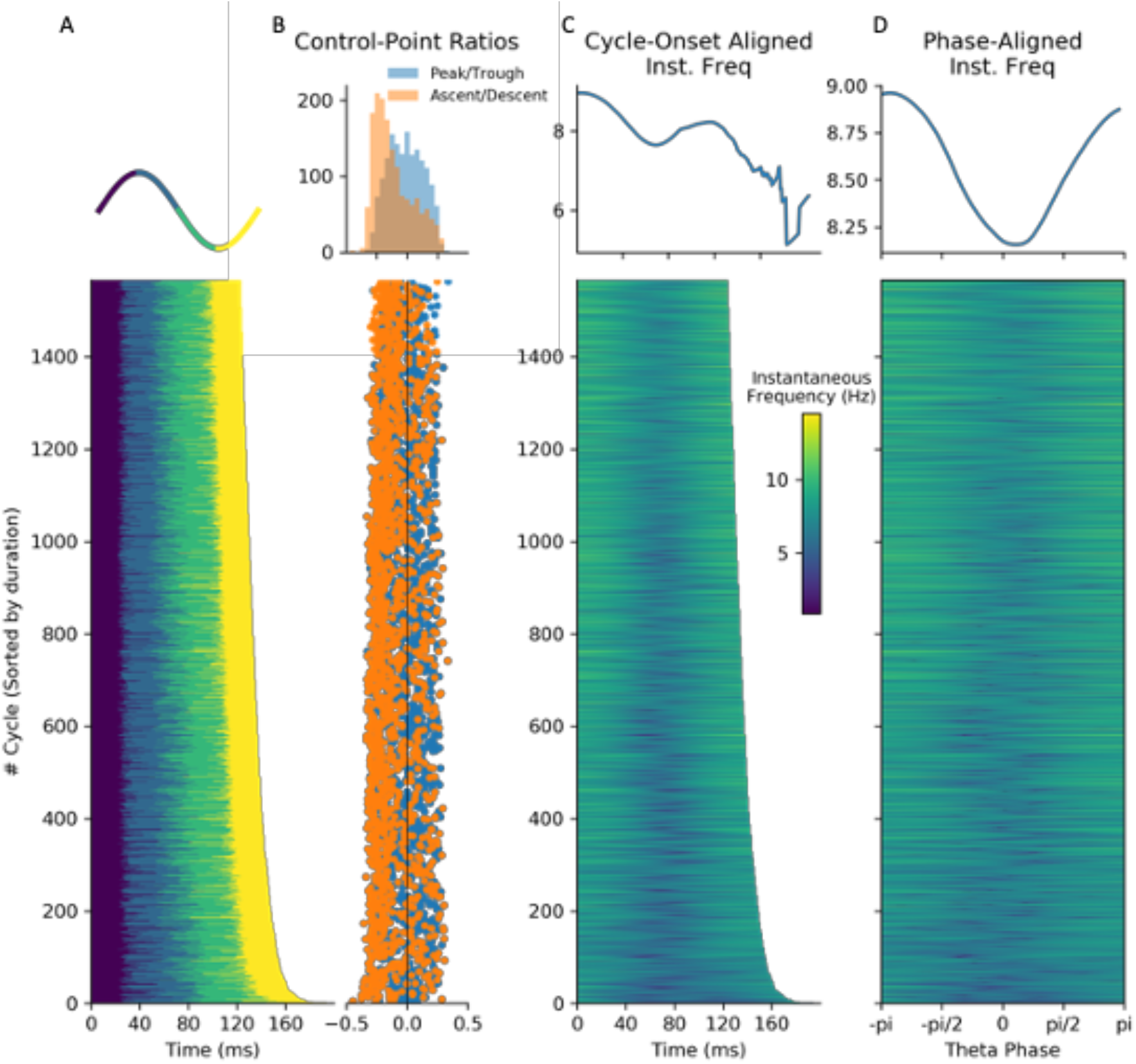
Methods for quantifying waveform in hippocampal theta. A : Durations between successive control points for each simulated cycle. B : Top: Distributions of peak-to-trough and ascent-to-descent durations. Bottom: Peak-to-trough and ascent-to-descent ratios for each cycle. C : Top: Average of temporally aligned instantaneous frequency profiles. Bottom: Temporally aligned instantaneous frequency profile for each cycle. D : Top: Average of phase-aligned instantaneous frequency profiles. Bottom: Phase-aligned instantaneous frequency profile for each cycle.

### 3.5: Theta has a stereotyped asymmetric shape with wide variability over cycles

We next summarised the average waveform across theta cycles from six recordings taken from three mice. The average phase-aligned instantaneous-frequency profile is computed for each recording and for the whole dataset. The overall group-level average waveform had a cosine-type profile centred around an average instantaneous frequency of ∼8.6 Hz (Figure 9A, average in black and individual recording sessions in grey). On average, the instantaneous frequency peaked within the cycle around 9 Hz at the ascending zero-crossing and drops to just below 8.4 Hz between the peak and descending zero-crossing. These results are consistent with previous studies showing an asymmetry between the fast-rising and slow-decaying halves of a theta cycle (Belluscio et al., 2012; Buzsáki et al., 1986; Cole and Voytek, 2019). All six recordings across three animals showed a shape with a maximum frequency around the ascending zero-crossing and a minimum on the descending edge, though there was some variability in whether the lowest frequency was closer to the peak or trough.

**Figure 9:**
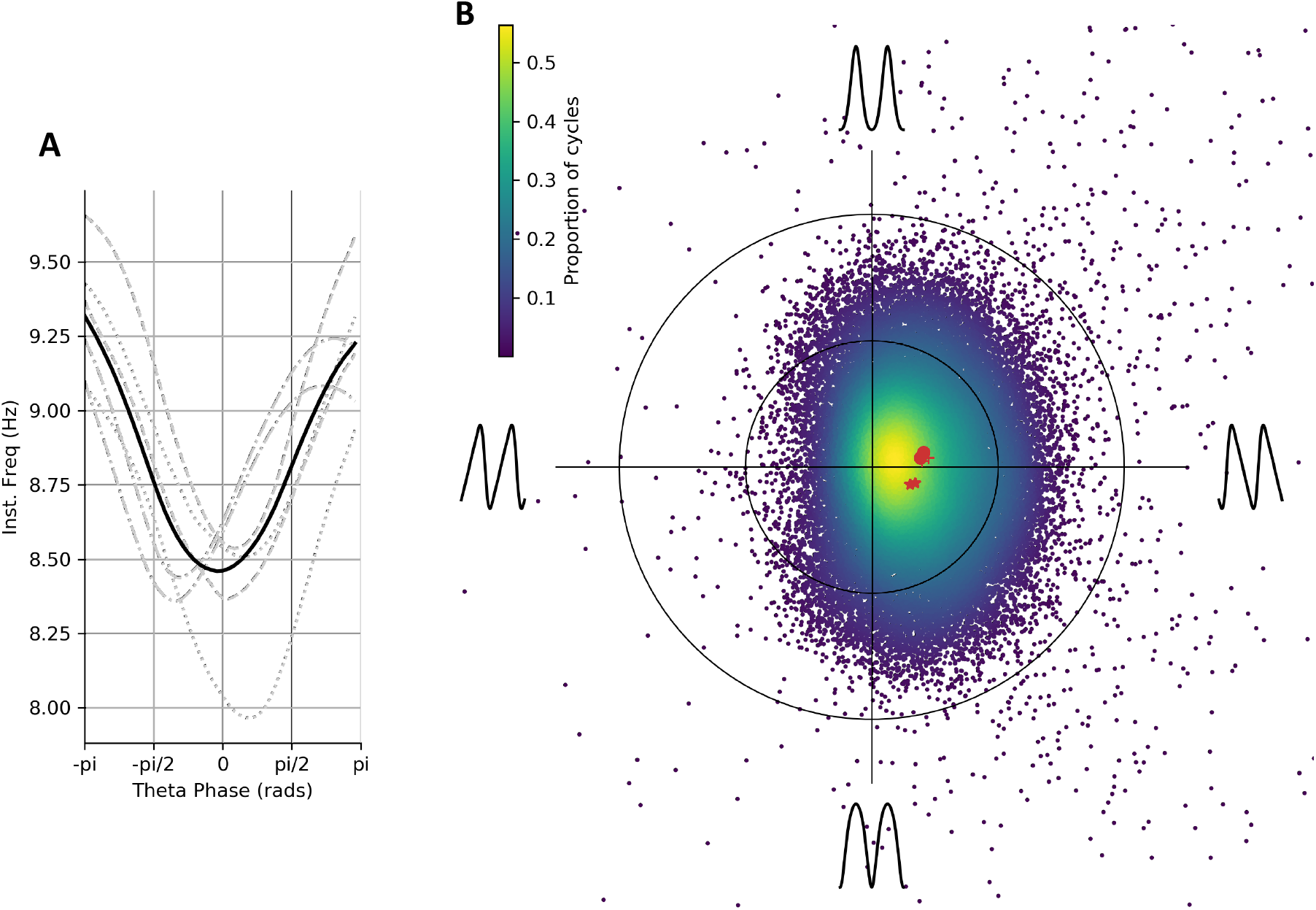
Average waveform shape and variability in shape across cycles in hippocampal theta. A : Average of phase-aligned instantaneous frequency profiles for each of the six separate recording sessions across three mice. The different dashed line styles indicate the runs from the different mice and the solid black line represents the average across all six recordings. B : Each individual cycle is projected into a simplified ‘shape-space’ to visualise the overall variability in waveform shape around the average. Individual recording averages are shown in red with different symbols representing the three animals.

To visualise the variability in waveform shape across cycles and recording sessions, we performed a complementary analysis using the instantaneous frequency mean-vector to see the distribution of single-cycle waveforms across a simplified 2-dimensional shape-space (Figure 9B; section 2.2.4). The distribution had a non-zero mean on the x-axis for all recordings, indicating that the highest frequencies in a cycle are typically at the ascending edge, consistent with the average in Figure 9A and with previous literature on the theta cycle (Belluscio et al., 2012). Though the overall mean shift in the distribution of cycles is robust, there is substantial cycle-to-cycle variability indicated by the width of the distribution.

### 3.6: Distinct waveform motifs are differentially related to behavioural and electrophysiological states

To further describe the variability in waveform shape across cycles and characterise its relation to movement speed, theta amplitude and theta cycle duration we identify a set of waveform shape ‘motifs’ using PCA. The PC values define a set of shape motifs that are each expressed to different degrees in the observed cycles (see section 2.2.5 and 2.4.4). The first four components describing 96% of variance are retained for further analysis (Figure 10). The waveform shape represented by each PC motif is summarised by the normalised waveforms (Figure 10A). These normalised waveforms are computed from the instantaneous frequency profiles of the PC component-vectors (Figure 10B) projected onto the extreme ends of the PC score distribution (Figure 10C). The shape of each individual cycle can then be described by a set of four PC scores, relating to the amount of each component which it contains.

**Figure 10:**
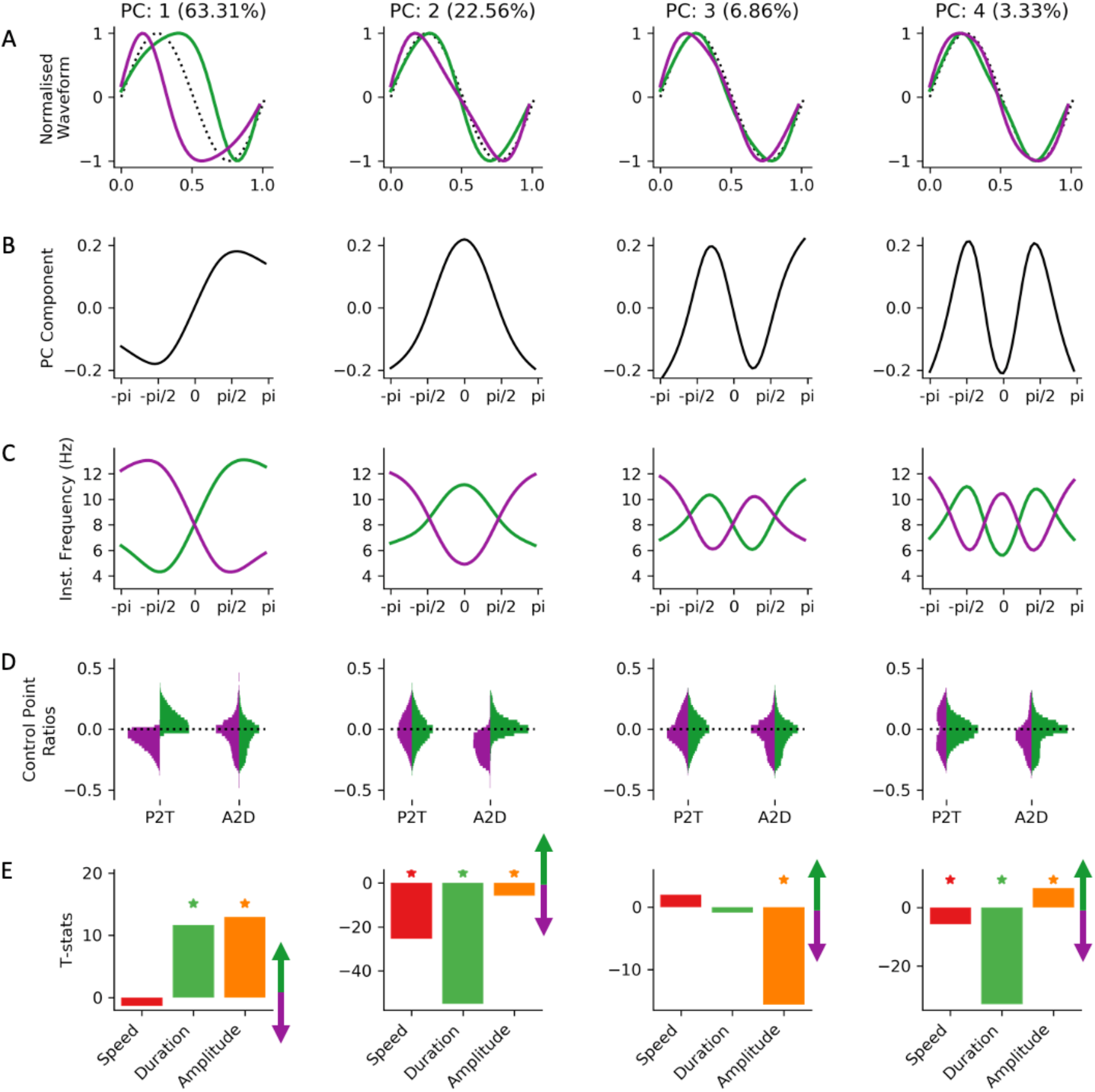
Shape motifs in hippocampal theta and their relation to movement speed. A : The normalised waveforms for the first four shape motifs identified from a PCA across all phase-aligned instantaneous frequency profiles. Waveforms for positive PC scores are shown in purple and waveforms for negative scores shown in green with a sinusoid for reference (black dotted line). B : PC for each motif. C : Instantaneous frequency profiles of each shape motif created by multiplying the PC shape in ‘B’ with the maximum or minimum observed PC score for that PC and adding the mean. Purple profiles represent the positive end of the score distribution and green profiles represent the negative end. D : Control-point ratios for cycles split by the sign of the PC score. Purple profiles represent the positive end of the score distribution and green profiles represent the negative end. E : t-value of a GLM modelling the PC score for each motif as a function of movement speed, theta cycle duration and theta cycle amplitude. Asterisks indicate statistical significance at p>0.01 as identified by non-parametric permutations.

PC-1 (63.31% of variance) describes a continuum of shape from a sharp peak and wide trough through to a wide peak and sharp trough. This shape is similar to the y-axis in the mean vector distribution in Figure 9B. In contrast, PC-2 (22.56% of variance) describes shapes ranging between an elongated ascending edge and an elongated descending edge, similar to the x-axis of Figure 9B. The remaining components describe more complex shapes with relatively small contributions to the variance explained. PC-3 (6.86% of variance) captures shapes with a left or right ‘tilt’ around their extrema and PC-4 (3.33% of variance) describes shapes with a sharper or flatter curvature around the extrema.

The control-point-based ascending-to-descending ratio and peak-to-trough ratio are computed for each cycle. For each PC, these values are partitioned into cycles with positive or negative PC scores (relating to distinct ends of the shape continuum for that component) and their distributions plotted in Figure 10D. The peak-to-trough ratios are clearly separated in the two ends of PC-1 whilst the ascending-to-descending ratios are similar for cycles with a positive or negative score in PC-1. This is consistent with the normalised waveforms summarising PC-1 in Figure 10A. PC-2 also shows the expected separation of ascending-to-descending ratios by PC score, whilst the peak-to-trough ratios are unchanged. Whilst PC-3 and PC-4 describe around 10% of shape variability, they are not characterised by the control point analyses. Neither peak-to-trough ratios nor ascending-to-descending ratios are changed by PC score for PC-3 or PC-4. These shape profiles are robustly identified by the phase-aligned instantaneous frequency method but are not distinguished by these control point-based metrics as the shape distortions in PC-3 and PC-4 occur between the four specified control points.

A general linear model was used to quantify the relationship between the different shape motifs and theta amplitude, theta duration and mouse movement speed. This regression is computed separately for each PC and the resulting parameter estimates converted into t-statistics. PC-1 codes for changes in average instantaneous frequency across the cycle with a small shape distortion around the descending edge. This PC has a strong relationship with cycle duration and amplitude but no significant covariation with movement speed. Longer and higher amplitude cycles tend to have more positive scores in PC-1 relating to wide peak shapes. PC-2 has a strong relationship with duration and movement speed. Specifically, cycles with elongated descending edges have longer cycle durations and are more likely to occur during fast animal movement. PC-3 shows significant covariance with cycle amplitude. High amplitude cycles tend to have shapes in which instantaneous frequency is relatively high just before the peak or trough. Finally, PC-4 varies strongly with duration and weakly with movement speed. Cycles with flatter curvatures around the extrema have longer cycle durations and are less likely to occur during faster animal movement.

## 4: Discussion

Non-sinusoidal waveforms are often visible by eye in raw LFP traces of electrophysiological datasets, yet discovering and quantifying these non-sinusoidal and non-linear features present substantial analytic challenges. We utilise within-cycle variability in instantaneous frequency to describe distortions in waveform (Huang et al., 2009). Further, we introduce phase-alignment as a solution to comparing full-resolution waveforms between cycles of different durations. Taken together, we establish that the phase-aligned instantaneous frequency profile of an oscillation provides a flexible framework for complete characterisation of oscillatory waveform shape. We demonstrate the utility of this approach by applying it to simulated data and LFP recordings of theta oscillations of behaving mice.

In real data, we observed that theta oscillations have, on average, a fast-ascending and slow descending waveform, in line with previous reports (Belluscio et al., 2012; Buzsáki et al., 1986, 1985; Cole and Voytek, 2019). Though this average shape is robust across many cycles, recording sessions and animals; the shape of individual cycles is highly variable. We characterise this variability using PCA to identify a range of shape components, or ‘shape motifs’, which maximally explain the variability in the dataset. The first two PCs quantify the relative durations of the peak and trough (PC-1) and the ascending to descending edge (PC-2). These PCs broadly map onto the features described by the peak-to-trough and ascending-to-descending control point ratios. We show that these theta shape PCs have distinct patterns of covariation with movement speed, theta amplitude and theta cycle duration. Critically, we show that though PC-2 describes less variability overall, it most clearly co-varies with movement speed.

PC-3 and PC-4 capture more complex waveform shapes. We show that the curvature around the extrema of the waveform shape (PC-4) is wider in theta cycles occurring during faster animal movement. This shape is naturally described by instantaneous frequency but not visible to standard ascending-to-descending and peak-to-trough control-point ratios. More generally, if the waveform shape of interest is known *a priori*, it is possible to construct specific control point-based measures so that the waveform shape can be identified. For instance, waveform sharpness can be explored by looking at the differential between the extrema and the samples 5ms before and after (Cole et al., 2017). However, in real data, we may not know the waveform shape of interest *a priori*, implying that many separate metrics may need to be computed for each cycle. In contrast, the phase-aligned instantaneous frequency can quantify any waveform shape as a within-cycle instantaneous frequency sweep without pre-specifying the features which may be of interest.

The present results demonstrate that single-cycle dynamics in oscillations can be meaningfully estimated using phase-aligned instantaneous frequency, and that specific shape motifs are differentially related to the wider electrophysiological (theta amplitude and duration) and behavioural (movement speed) context. Future models of theta function may consider these dynamics in waveform shape that deviate from a canonical sinusoidal theta template. Given that many sub-processes occur preferentially at different parts of the theta cycle (Klausberger and Somogyi, 2008), we hypothesise that shape distortion may indicate or reflect a change in the underlying theta-phase-nested sub-processes.

The outlined approach requires that each cycle is smooth in both its waveform and phase profiles, as any jumps or discontinuities will lead to noisy or even negative instantaneous frequency estimates. If the cycle is smooth, we can characterise very large distortions in waveform shape as within-cycle dynamics in instantaneous frequency. Finally, we assume that the features being analysed are well described as oscillations. If the features are non-sinusoidal and non-oscillatory, such as spiking activity, then descriptions using the language of frequency may not be appropriate. With these improvements and caveats in hand, this approach is readily generalisable to other datasets and provides a flexible framework for investigating waveform shape oscillating systems.

In conclusion, the full-cycle waveform of single cycles of hippocampal theta can be quantified and explored with phase-aligned instantaneous frequency. We use this approach to confirm the characteristic fast-ascending waveform of theta oscillations; and to additionally reveal that this is highly variable on the single-cycle level. Moreover, we are able to link this variability with behavioural and electrophysiological states, suggesting that waveform shape is a relevant feature of neuronal oscillations alongside frequency, phase and amplitude. Finally, whilst we have illustrated this approach with hippocampal theta oscillations, it is likely that this methodology will readily generalise to neuronal oscillation in other brain regions, frequency bands and contexts.

## 5: Acknowledgments & Funding

The authors would like to thank Catharina Zich for discussion of the manuscript. This project was supported by the Medical Research Council (RG94383/RG89702) and by the NIHR Oxford Health Biomedical Research Centre. The Wellcome Centre for Integrative Neuroimaging is supported by core funding from the Wellcome Trust (203139/Z/16/Z). V.L.d.S. and D.D. are supported by the Medical Research Council UK (Programmes MC_UU_12024/3 and MC_UU_00003/4 to D.D.) ACN is supported by the Wellcome Trust (104571/Z/14/Z) and James S. McDonnell foundation (220020448). MWW is supported by the Wellcome Trust (106183/Z/14/Z; 215573/Z/19/Z). ACN and MWW are further supported by an EU European Training Network grant (euSSN; 860563).

This research was funded in whole, or in part, by the Wellcome Trust. For the purpose of Open Access, the author has applied a CC-BY public copyright licence to any Author Accepted Manuscript version arising from this submission.

## 6: Author contributions

Conceptualisation: AJQ, VLdS, NH, CHJ, WKL, JRY, ACN, DD and MWW.

Methodology: AJQ, VLdS, NH, CHJ, WKL, JRY, ACN, DD and MWW.

Resources: VLdS, ACN, DD and MWW.

Investigation: AJQ, VLdS, DD and MWW.

Formal analysis: AJQ.

Software: AJQ.

Visualisation: AJQ.

Writing Original Draft: AJQ, VLdS, ACN, DD and MWW.

Writing Review & Edit: AJQ, VLdS, NH, CHJ, WKL, JRY, ACN, DD and MWW.

Supervision: ACN and MWW.

Funding Acquisition: DD, ACN and MWW

## 8: Supplemental Methods

### Experimental model and subject details

Animals used were male adult (4–7 months old) C57BL/6J mice (Charles River, UK). All animals had free access to water and food in a dedicated housing facility with a 12/12 h light/dark cycle. They shared a cage with their littermates until the surgery. All experiments involving animals were conducted according to the UK Animals (Scientific Procedures) Act 1986 under personal and project licenses issued by the Home Office following ethical review.

#### Microdrive implantation

Animals were implanted with a 10-12 tetrode microdrive during a surgical procedure performed under deep anesthesia using isoflurane (0.5–2 %) and oxygen (2 l/min), with analgesia (0.1 mg/kg vetergesic) provided before and after. Tetrodes were constructed by twisting together four insulated tungsten wires (12 µm diameter, California Fine Wire) and shortly heating them to bind them together in a single bundle. Each tetrode was attached to a M1.0 screw to enable their independent movement. The drive was implanted under stereotaxic control in reference to bregma (Lopes-dos-Santos et al., 2018). Tetrodes were initially implanted above the CA1 pyramidal layer and their exposed parts were covered with paraffin wax. The drive was then secured to the skull using dental cement. For extra stability, stainless-steel anchor screws had first been inserted into the skull. Two of the anchor screws, which were inserted above the cerebellum, were attached to 50 µm tungsten wires (California Fine Wire) and served as ground and reference electrodes during the recordings. The placement of the tetrodes in dorsal CA1 was confirmed by the electrophysiological profile of the local field potentials in the hippocampal ripple frequency band.

#### Recording procedures

Recordings commenced following full recovery from the surgery. Each animal was connected to the recording apparatus and familiarized with a high-walled box containing home cage bedding and with one open-field enclosure (the familiar enclosure) over a period of approximately seven days. During this period, tetrodes were gradually lowered to the stratum oriens of the hippocampal CA1. On the morning of each recording day, tetrodes were further lowered into the pyramidal cell layer in search of multi-unit spiking activity and sharp-wave/ripple events (van de Ven et al., 2016). Tetrodes were not moved for at least 1.5 h before recordings started. For each recording day, the animal was exposed to various open-field enclosures including the familiar, which the animal had repeatedly been exposed to before, and a novel enclosure the animal had never seen before. The open-field enclosures differed in shape and in the cue-cards that lined some of the walls. The present study includes a total of six LFP recordings from three mice (including three familiar enclosure and three novel enclosure sessions). At the end of each recording day, tetrodes were raised to the stratum oriens to avoid damaging the pyramidal layer overnight.

#### Multichannel data acquisition and position tracking

The extracellular signals from the electrodes were buffered on the head of the animal (unity gain op-amps, Axona Ltd) and transmitted over a single strand of litz wire to a dual stage amplifier and band pass filter (gain 1000, pass band 0.1 Hz to 5 kHz; Sensorium Inc., Charlotte, VT), or (in other setups) the electrode signals were amplified, multiplexed, and digitized using a single integrated circuit located on the head of the animal (RHD2164, Intan Technologies, Los Angeles; pass band 0.09 Hz to 7.60 kHz). The amplified and filtered electrophysiological signals were digitized at 20 kHz and saved to disk along with the synchronization signals from the position tracking. LFPs were further down sampled to 1250 Hz for all subsequent analyses. In order to track the location of the animal three LED clusters were attached to the electrode casing and captured at 39 frames per second by an overhead color camera.

### 8.2: Mask sift parameter robustness

We ran a supplemental analysis to ensure that our LFP phase-aligned instantaneous frequency results were robust to moderate changes in the masking parameters. 5 minutes of data from a single data recording were repeatedly sifted with jittered masked frequencies. 25 sifts were computed for each of three mask frequency jitter values of 10%, 20% and 30%. For example, in the 10% condition each iteration each mask frequency was randomised to a value drawn from a uniform distribution between 90% and 110% of the original frequency. The results showed that jitter of 10% has a small effect on the phase-aligned instantaneous frequency values, though the centre frequency and shape profile remain consistent across all iterations. Jitters of 20% and 30% have larger effects on both centre frequency and shape on individual iterations, suggesting that some mask frequency combinations are having a large impact on the results. Despite this, the average across all iterations remains strikingly similar. These results indicate that the main waveform results are robust to moderate changes to the masking parameters.

**Figure S1.**
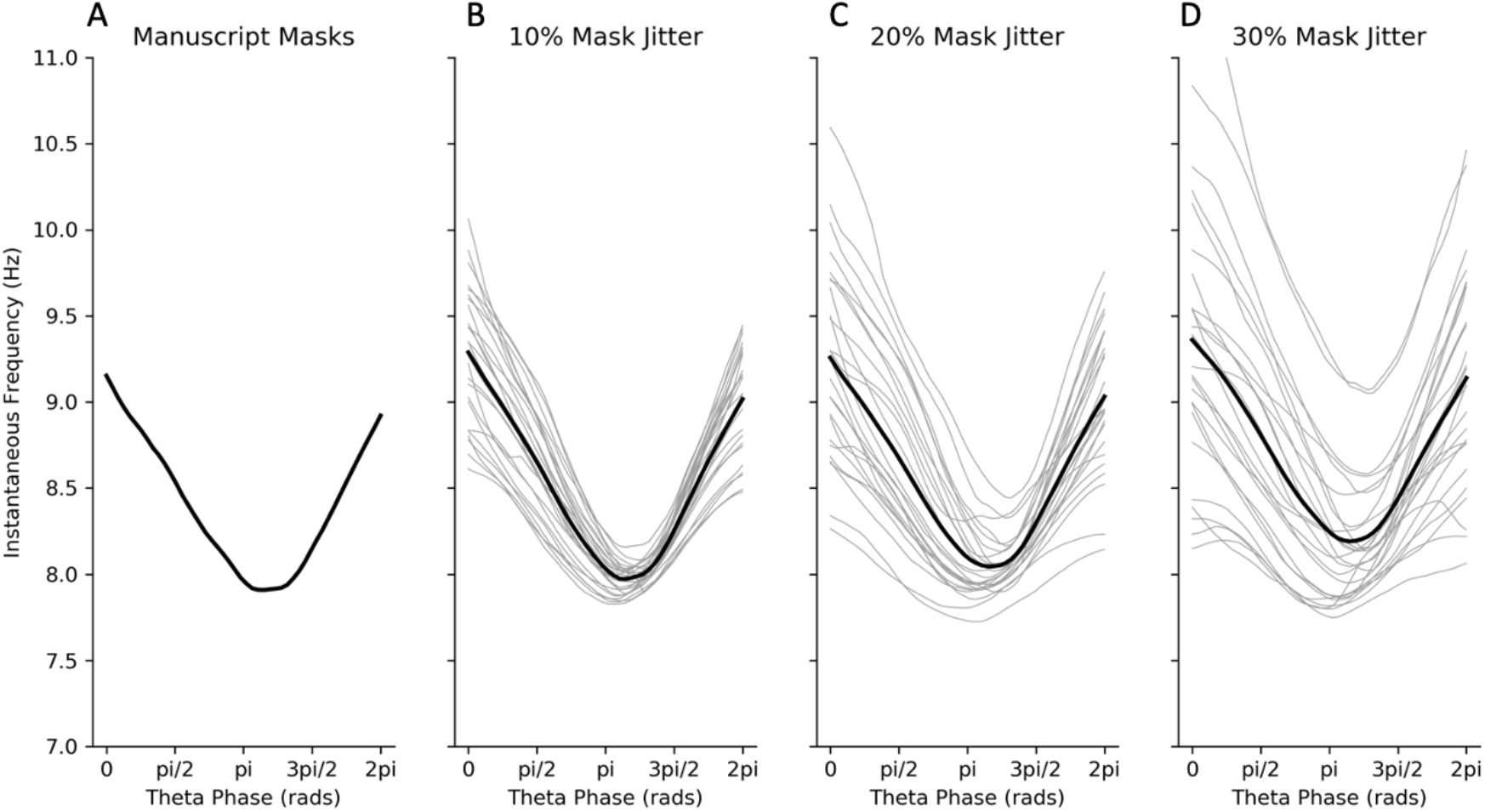
Phase-aligned instantaneous frequency values across a range of jittered mask frequencies. A: Instantaneous frequency for mask frequencies used in main analysis B: Instantaneous frequency for mask frequencies jittered by +/-10%. Individual iterations are shown in grey and the average in black. C: As B for jitter of +/-20% D: As B for jitter of +/-30%

### 8.3 : PCA component selection and reproducibility

The principal components analysis results were validated by computing the split half reproducibility of the PC components across 500 splits. The correlation of component shapes between the separate halves and the proportion of variance explained was computed for each split. The distribution of explained variance for each mode was highly reproducible across the 500 splits and the first four components explained more than 5% of overall variance (Figure S2A). The PC component shapes were also highly reliable for the first four components. The average correlation between components for the two halves of each split was over r=0.95 for the first 5 PCs. Based on these comparisons we carried the first four components forward for further analyses.

**Figure S2.**
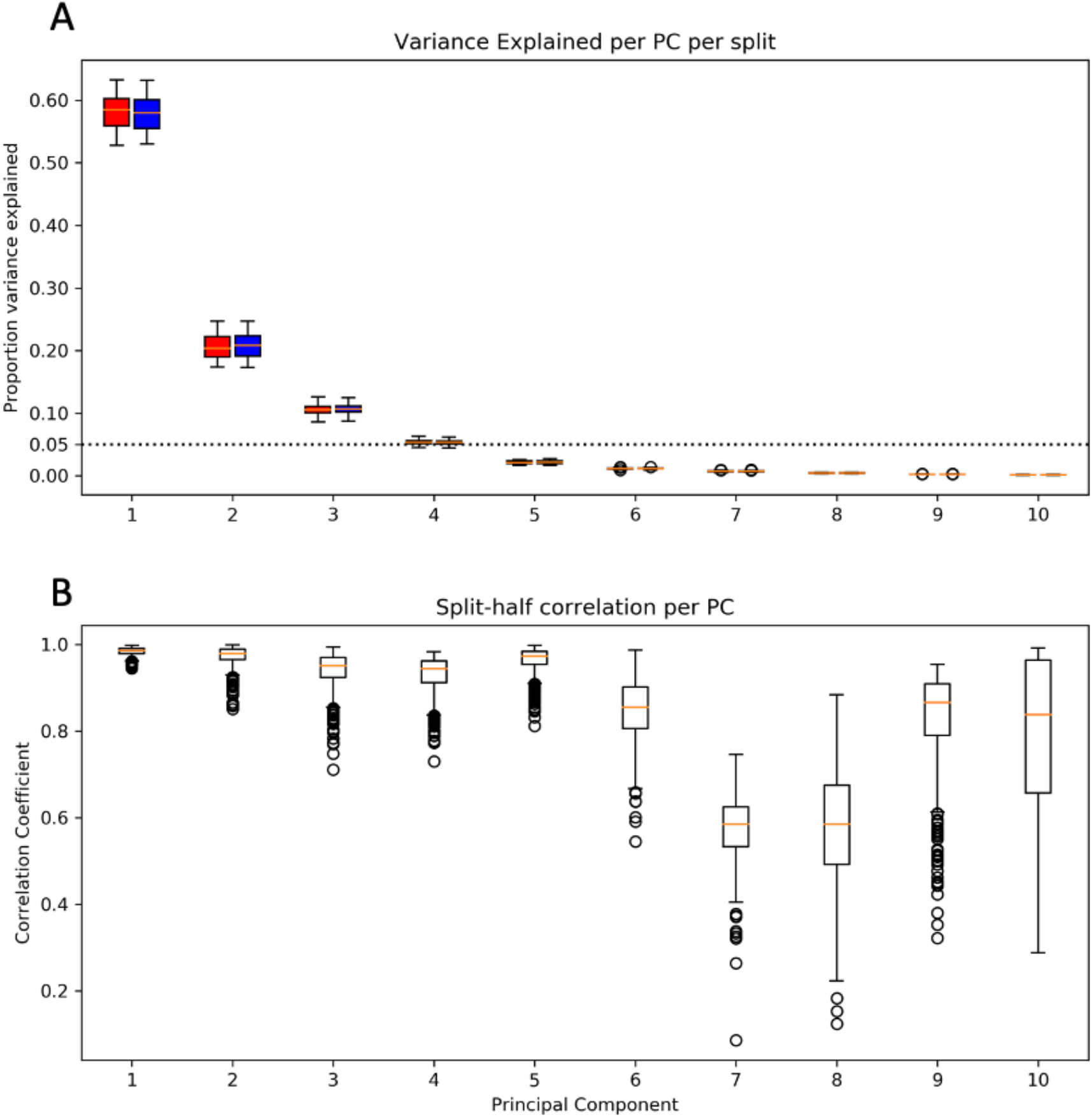
the variance explained and split-half correlation distributions across 500 split half iterations. A: the variance explained by each component for each half over the 500 splits. The first half is in red and the second half in blue. B: The correlation in the component shape between the first and second half of the 500 splits.

